# Controlled and polygynous mating in the black soldier fly: advancing breeding programs utilizing quantitative genetic designs

**DOI:** 10.1101/2024.09.09.611978

**Authors:** K. Jensen, S.F. Thormose, N.K. Noer, T.M. Schou, M. Kargo, A. Gligorescu, J.V. Nørgaard, L.S. Hansen, R.M. Zaalberg, H.M. Nielsen, T.N. Kristensen

## Abstract

In animal breeding programs, utilizing quantitative genetic designs such as the full-/half-sibling design is fundamental. A full-/half-sibling design demands that mating can be controlled, and individuals can be tracked, enabling construction of a pedigree. In nature, black soldier fly (*Hermetia illucens* (L.)) males are reported to gather in lekking groups to engage in competitive displays and courtship rituals before mating, and this lekking behavior is described as crucial for mating in this species. We show that a randomly chosen male and a randomly chosen virgin female can readily mate under LED lights in a small container at 28 °C and 70% RH, demonstrating the ability to mate selected pairs. For males given access to four females, all mated and the time until first mating was considerably shorter than for males given only one female, where 80% of the males mated. Furthermore, when adding four virgin females to a single male, males successfully mated all four females in 52% of cases within our four-hour trial, with 96% of males mating at least twice. Number of matings observed for a male correlated with the number of females from the container that laid an egg clutch, and almost all egg clutches produced offspring. Observed mating thus lead to fertilization, and consecutive matings were with new females resulting in full- and half-sibling offspring. Males with four females that mated a second female did so at a median time of less than three minutes upon ending previous mating, showing that polygynous mating occurred rapidly in this setup. Our findings pave the way for moving *H. illucens* breeding programs beyond mass selection towards advanced selective breeding designs where controlled mating is typically required. Such designs enable selection for multiple traits simultaneously while controlling inbreeding and can drastically increase rates of selection responses compared to mass selection.

## Introduction

Insects are currently being proposed as a sustainable future protein and lipid source (Dobermann *et al*., 2017; Liceaga, 2021; van Huis, 2020), and some insect species have been used as food and feed for centuries (Olivadese and Dindo, 2023). In the Western world, large-scale factories producing insect-derived products have only recently been established (Montanari *et al*., 2021). Especially species of flies, beetles, and crickets are target taxonomic groups gaining attention (Oonincx *et al*., 2015; van Huis, 2021). These species differ from traditional livestock in many aspects of biology and management. Since the domestication of most insect species began recently and is still in progress, there is much to be learned about their biology, adaptation, and performance under artificial rearing conditions. One prerequisite for efficient selective breeding is the ability to perform controlled matings of selected pairs, but some insect species may have specific requirements to engage in mating (Burdfield-Steel and Harari, 2021). Knowing how to mate selected pairs is essential when developing breeding programs aimed at improving traits of interest in commercial insect species through selective breeding (Hansen *et al*., 2024b).

Selection has been instrumental in securing continued and cumulative genetic progress for production traits in livestock for decades (Hill and Kirkpatrick, 2010). Genetic improvement of livestock and their feed, together with feeding optimization and technological advancements, plays a key role in producing sustainable food and feed for a growing human population. In traditional livestock such as cattle, pigs, and poultry, the effectiveness of breeding programs has relied heavily on the ability to master the reproduction of the animals. This has resulted in more profitable animals that e.g. utilize feed for meat growth at a much higher efficiency (Emmerson, 1997; Lashley, 1978; Lush, 1937; Schultz *et al*., 2020). Another reason for the effectiveness of livestock selective breeding is the ability to keep track of pedigrees across generations, which greatly improves the precision by which genetically superior animals can be identified and thus boosts breeding responses (Clark *et al*., 2012; Meuwissen, 1997). Not only does this allow for fast genetic progress across generations but also for control of the rate of inbreeding in a population (Meuwissen, 1997), as well as for effective selection on multiple traits simultaneously (Cole *et al*., 2021).

Genetic improvement of insect species for food and feed is still in its infancy (Gowda *et al*., 2025; Hansen *et al*., 2024b; Jensen *et al*., 2017). Yet, the potential for fast genetic improvement through selective breeding programs in insects is high given that there is plenty of genetic variation within relevant species (Cai *et al*., 2024; Eriksson and Picard, 2021; Hoffmann *et al*., 2021), most insects have short generation intervals spanning from weeks to a few months, and the small size of individuals means that high census population sizes can be maintained on relatively little space. However, to utilize this potential we need to be able to control mating to secure that selected individuals are parents of the next generation, excluding any chance for polyandry.

Recent studies aiming to genetically improve production traits in commercially produced insects have done so employing mass selection (Facchini *et al*., 2022; Gligorescu *et al*., 2023; Morales-Ramos *et al*., 2019). However, under typical mass selection conditions no information is available about exactly which flies mate and contribute genetically to the next generation, and exactly which eggs are from which parents. The pedigree structure is therefore unknown. Pedigree information can facilitate the estimation of trait heritabilities, variance components, and genetic correlations between traits of interest (Falconer and Mackay, 1996). One commonly used method for estimating such genetic parameters is through a full-/half-sibling design, where phenotypes of siblings and half-siblings are compared. Many insect species will readily mate when placed together as virgins in pairs, especially amongst insects typically maintained in laboratory cultures, including crickets (Hawkes *et al*., 2022; Rapkin *et al*., 2017), cockroaches (Bunning *et al*., 2015; Jensen and Silverman, 2018), silkworms (Abdelmegeed, 2015; Sarkar *et al*., 2009), mealworms (Sellem *et al*., 2024; Worden and Parker, 2001), fruit flies (Jensen *et al*., 2015; Perez-Staples *et al*., 2007), and house flies (Hansen *et al*., 2024a; Laursen *et al*., 2024). Full-/half-sibling designs have been successfully applied in the house fly (Boatta *et al*., 2023; Hansen *et al*., 2024a), and full-sibling designs have been applied in mealworms (Sellem *et al*., 2024). However, many insect species require specific conditions to induce mating (Burdfield-Steel and Harari, 2021; Lloyd, 1979), and mating selected individuals is therefore not easily attainable for all species.

The black soldier fly (*Hermetia illucens* (L.); Diptera: Stratiomyidae) has become a main species of interest in the insect production industry (Tomberlin and van Huis, 2020; van Huis *et al*., 2020), and there is great interest in genetically improving this species for production through breeding (Athanassiou *et al*., 2025). Studies on *H. illucens* have shown that traits related to developmental time, feed conversion efficiency, and weight gain can be improved under mass selection (Facchini *et al*., 2022, Gligorescu *et al*., 2023). However, this type of selection does not easily allow for controlling rates of inbreeding, for selecting on multiple traits simultaneously, and long-term responses to selection are expected to be lower than what can be obtained when pedigree information is available (Falconer and Mackay, 1996). In nature, *H. illucens* have lekking behavior (Barrett *et al*., 2023; Chiabotto *et al*., 2024; Giunti *et al*., 2018; Jones and Tomberlin, 2021; Julita *et al*., 2020; Kortsmit *et al*., 2023; Lemke *et al*., 2023; Meneguz *et al*., 2023; Muraro *et al*., 2024; Tomberlin and Sheppard, 2001). Lekking occurs when several males perform sexual display together to attract females while each male defends a territorial patch (Alcock, 1990; Fiske *et al*., 1998; Tomberlin and Sheppard, 2001), and it has been a general belief that a single male is unable to attain successful courtship (Barrett *et al*., 2023; Chiabotto *et al*., 2024; Giunti *et al*., 2018; Jones and Tomberlin, 2021; Julita *et al*., 2020; Kortsmit *et al*., 2023; Laudani *et al*., 2024; Lemke *et al*., 2023; Manas *et al*., 2024; Meneguz *et al*., 2023; Muraro *et al*., 2024; Tomberlin and Sheppard, 2001). If this is the case, controlled mating where a specific male is mated to a specific female is difficult to perform. Using marked flies, keeping track of mated pairs is possible (Chiabotto *et al*., 2024; Jones and Tomberlin, 2021; Muraro *et al*., 2024), but clearly tedious. If the presence of multiple males is necessary to induce mating in *H. illucens* as reported to this point, this is a major challenge for developing effective selective breeding programs involving pedigree tracking and full-/half-sibling designs for this species.

Optimal conditions for mating to occur in *H. illucens* include moderately high temperatures between 26 to 29 °C and a high relative humidity of 65 to 85 % (Giunti *et al*., 2018; Julita *et al*., 2020). Bright light with high intensity in the blue-green spectrum is critical for encouraging mating (Barrett *et al*., 2023; Heussler *et al*., 2018; Liu *et al*., 2020; Oonincx *et al*., 2016; Schneider, 2020; Tomberlin and Sheppard, 2002), and LED lights have proven to be the best artificial light source in comparative tests (Heussler *et al*., 2018; Hoc *et al*., 2019; Julita *et al*., 2020; Liu *et al*., 2020; Macavei *et al*., 2020; Oonincx *et al*., 2016). Although it has been demonstrated that both male and female *H. illucens* can mate more than once (Chiabotto *et al*., 2024; Dickerson *et al*., 2024; Hoffmann *et al*., 2021; Jones and Tomberlin, 2021; Muraro *et al*., 2024), little is known about how quickly *H. illucens* males will mate polygynously when multiple females are present. If controlled polygynous mating of individual males is possible, this would enable the development of efficient pedigree-based breeding programs for this species (Slagboom *et al*., 2024). Furthermore, controlled polygynous mating would create opportunities to advance selective breeding strategies, for example by simultaneously selecting for multiple traits of economic value (Zaalberg *et al*., 2024). Our primary goals were to investigate whether males and females can mate in individual pairs within small containers with no chance for partner choice, and whether and how quickly individually maintained males can mate multiple virgin females consecutively. We then investigated whether mating led to egg clutches and offspring, and whether the number of offspring produced by a female was affected by the total number of females the male had mated. Our findings provide a foundation for implementing controlled mating in *H. illucens* breeding programs, contributing to the advancement of pedigree-based selective breeding in commercial insect production.

## Materials and methods

### Fly population and rearing

We used a laboratory culture of *H. illucens* established in 2017 at Enorm Biofactory A/S (Hedelundvej 15, 8762 Flemming, Denmark) from a batch of >100,000 juveniles. The culture was maintained in a large population of >5,000 adults per generation prior to this experiment. Rearing of flies and all experiments were performed in climate-controlled rooms kept at 28 ± 0.5 °C and 70 ± 10% relative humidity (RH). Larvae were collected from Enorm at 5-6 days of age and reared in open trays (57:37:8 cm L:W:H - internal measures) each containing two kg of chicken feed (Paco Start, Danish Agricultural Grocery Company, Fredericia, Denmark) and 4 L of water (66.67% substrate humidity by mass). Roughly 1500 larvae (15 g) were added to each tray. To control fly age since eclosion and ensure virginity (Permana *et al*., 2020), pupae were collected at their final stage and maintained in individual drosophila vials (95:25 mm height:diameter; VWR) secured with a foam stopper, and checked daily for emergence. Emerged adults were then given another 48-72 h in their individual vials to sexually mature. They were then sexed in the vial by visual inspection of the genitalia (Julita *et al*., 2020) and used in mating trials. All flies in mating trials thus were of a similar age (48-72 h), ensuring virility and sexual maturity (Dickerson *et al*., 2024; Tomberlin and Sheppard, 2002). To test for occurrence of unfertilized oviposition, a parallel set of 247 female flies were maintained in their individual vials since collection as pupae and monitored for the appearance of egg clutches until their death.

### Experimental design

For the mating experiments, male and female flies were maintained together in transparent 3.0 L (19.5:19.5:11.3 cm L:W:H) plastic containers (TPS192, Dansk Transport Emballage A/S, Vojens, Denmark). An eight cm diameter hole was cut in the lid for ventilation and covered with a fine mesh. All flies were allocated to the containers by simultaneously removing all foam stoppers and shaking flies out of their vials into the containers through an opened corner of the lid. This was done in a room at 20 °C, 50% RH, and lighting from fluorescent tubes in the roof, which stimulated very limited activity and no mating display. The containers with flies were then moved to the climate-controlled room for mating trials.

We established 30 replicates of one male with one female, and 25 replicates of one male with four females. All flies had an age range of 48-96 h since eclosion when set up in trial. Here, the flies were given four hours to mate under two LED lights (112.5 cm 50W Philips LED*life* VerticalGrow120 full spectrum) placed 8 cm above the container lids. All replicates involving one male with one female were performed on one day, and the replicates involving one male with four females were performed over the following two days with ten replicates on the first day and 15 on the second. Number of trials on a day fitted with the observation capacity of a single observer. We recorded the time until start of each mating by manual observation and recording using a timer and notebook. For the males given access to four females, we also measured time until the end of each mating. The timer was started when all flies were placed in the climate-controlled room underneath the LED lights, and the LED lights were turned on. No mating related activity occurred before this point. After the four hours with the male in the container, females were caught and placed in individual drosophila vials held upside down with coarse polyurethane foam stoppers (SuperFish aquarium filter, one-two mm spaces) clogging the bottoms for ovipositing. To enhance egg survival, a tray (57:37:8 cm L:W:H) with water was placed 10 cm underneath the vials, ensuring that RH did not drop below 70%, and vials were checked daily for egg clutches over the following six days. All egg clutches were placed in individual drosophila vials by collecting the foam stopper with the eggs and plugging it deep into the vial, so the eggs were about one cm from the surface of a two cm layer substrate of 5 g chicken feed with 10 mL water. After seven days, the vials with larvae were recorded and larvae with remaining substrate were transferred to an eight cm diameter plastic container with 30 g chicken feed and 60 mL water and given seven more days to grow. The container was covered with a lid with a six cm diameter hole with a fine mesh for ventilation. The larvae were then collected from their media and counted.

### Statistical analysis

Homogeneity of variances were compared between groups with a Levene’s test, and Shapiro-Wilk tests were used to test for normal distribution of residuals. Variances were in most cases unequal (Levene tests, *P* < 0.05), and data were generally not normally distributed (Shapiro– Wilk tests, *P* < 0.05). Therefore, nonparametric tests were used. A Kruskal-Wallis test was used when comparing time until each consecutive mating since start or since the end of the previous mating, as well as when comparing mating duration across consecutive matings. It was also used when testing for an effect of the number of times a male mated on the number of larvae produced per egg clutch by females from the same container. If the Kruskal-Wallis test indicated significant differences, it was followed by a Dunn’s test for multiple comparisons. A Wilcoxon test was used to compare the time until first mating and the number of larvae produced per female egg clutch by males paired with one *versus* four females. A Pearson’s chi-squared test was used to test for a difference in the proportion of males that mated within the four-hour mating trial depending on whether they were exposed to one or four females. This test was also used to test for a correlation between the number of times a male was observed mating and the number of females that laid an egg clutch, and to test for an effect of multiple mating on the proportion of egg clutches that resulted in larvae within males exposed to four females. Statistical analyses were performed in JMP 15.0.0 (SAS Institute Inc., Cary, NC, USA).

## Results

### Time until first mating

All single males exposed to four females mated at least once, while 80% of single males provided access to only a single female mated, which was a significantly lower proportion (Pearson: □^2^_1,55_ = 5.61, *P* = 0.018; Figure 1). Most males exposed to four females also mated a second (96%) and a third time (88%), and just above half of the males (52%) mated four times. The time to first mating was considerably shorter when males were together with four females than when they were together with only one female (Figure 1). Across the 25 males provided access to four females, the median time until first mating was 07:17 (mm:ss) minutes (10^th^ percentile: 03:46, 25^th^: 05:04, 75^th^: 10:46, 90^th^: 21:38), while males exposed to only one female initiated mating at a median of 53:58 minutes (10^th^ percentile: 07:40, 25^th^: 13:29, 75^th^: 01:04:24 (hh:mm:ss), 90^th^: 02:50:00). All males exposed to four females had initiated mating within 35 minutes (Figure 1).

**FIGURE 1.**
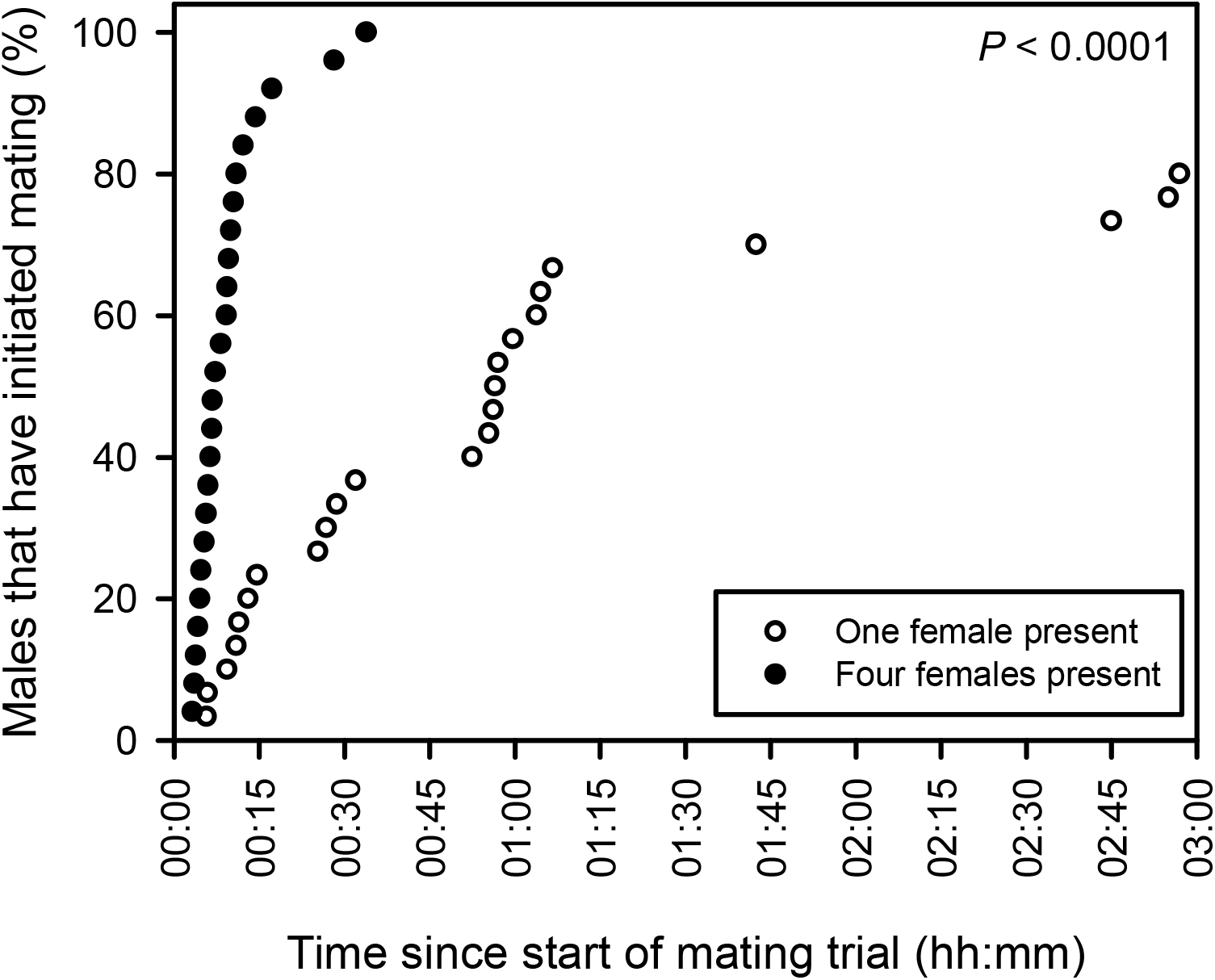
Timepoints and cumulative percentages of males initiating their first mating when given access to one (white) or four (black) females. The *P*-value is from a Wilcoxon test on time until mating depending on the number of females present, including only males that mated.

### Time between matings and mating durations

Males given four females that mated a second time initiated their second mating at a median of 02:43 minutes (10^th^ percentile: 00:48, 25^th^: 01:28, 75^th^: 08:11, 90^th^: 10:44) after the first mating had ended, and all had initiated their second mating within 30 minutes after terminating the first (Figure 2A). After completing the second mating, the median time until next mating increased significantly (Figure 2A). However, a minimum of 10 seconds was recorded for two males to initiate the third mating after finishing the second. Mating duration generally increased for each mating from around 20 minutes for the first mating to around 40 minutes for the third, with an intermediate 30 minutes of mating for the fourth mating (Figure 2B).

**FIGURE 2.**
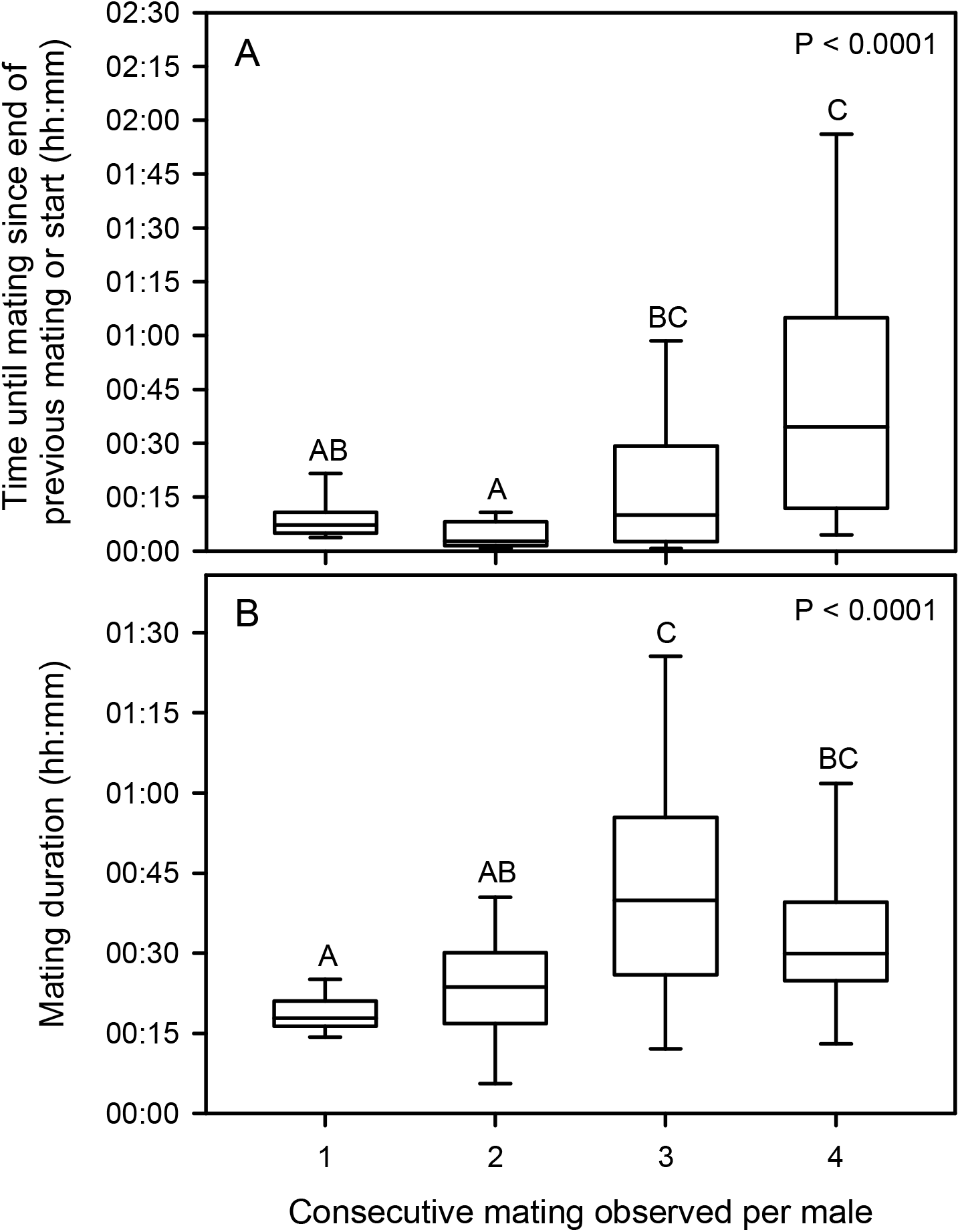
A) Time until first mating and in between subsequent matings and B) duration of each consecutive mating in males exposed to four virgin females for four hours. Boxes show median and 10^th^, 25^th^, 75^th^, and 90^th^ percentiles. Outliers are not shown. The *P*-values are from Kruskal-Wallis tests across the consecutive matings. Different letters indicate significant differences (Dunn test, α = 0.05).

### Egg clutch and offspring production

Out of the 30 females set up individually with a male, 23 subsequently produced an egg clutch, and all egg clutches produced larvae. A median of 149 larvae (10^th^ percentile: 16, 25^th^: 79, 75^th^: 210, 90^th^: 310) were produced from these egg clutches. For the 25 males with access to four females, in 8% (2) of the cases only one female produced an egg clutch, in 44% (11) of the cases two females produced an egg clutch, in 36% (9) of the cases three females produced an egg clutch, and in 12% (3) of the cases all four females produced an egg clutch.

We found a significant correlation between the number of times a male was observed mating and the number of females from the container that produced an egg clutch (Pearson: □^2^_9,25_ = 18.32, *P* = 0.032). Within these 63 egg clutches, 92% (58) produced larvae. This demonstrates that most matings lead to fertilization, and that matings were predominantly with different, virgin females. In the 247 female flies maintained in their individual vials since collection as pupae, 4% (10) laid an egg clutch within their lifetime, confirming that little oviposition occurred without prior mating and fertilization.

### Effects of multiple mating on fertility

The proportion of egg clutches that produced offspring was not affected by the number of times a male was observed mating (Pearson: □^2^_3,63_ = 4.73, *P* = 0.19). The egg clutches each produced a median of 138 larvae (10^th^ percentile: 33, 25^th^: 69, 75^th^: 219, 90^th^: 269), with no effect of the number of females from the container that oviposited (Kruskal-Wallis: □^2^_3,58_ = 3.41, *P* = 0.33), and there was similarly no significant difference in the number of larvae produced per egg clutch depending on whether one or four females were available to a male (Wilcoxon: □^2^_1,81_= 0.007, *P* = 0.93). These findings indicate that no sperm depletion occurred.

## Discussion

The importance of adult behavior is increasingly recognized in the breeding and maintenance of *H. illucens* populations (Lemke *et al*., 2023; Meneguz *et al*., 2023). With this study, we aimed to answer questions of central importance for the ability to execute efficient selective breeding programs in *H. illucens*. The main aims were to investigate whether *H. illucens* males and females can mate in individual pairs under controlled conditions with no chance for partner choice, and whether individually maintained *H. illucens* males can consecutively mate multiple virgin females within a short mating session and produce full- and half-sibling offspring with these. Our results show that *H. illucens* males and females can mate readily in controlled mating pairs of selected individuals without prior lekking within our setup. In effect, we have heavily relaxed sexual selection, allowing for predominantly artificial selection which is optimal in selective breeding. We furthermore show that multiple successful matings can occur quickly between one male and several females within our setup, showing that *H. illucens* can be highly polygynous and can readily produce full- and half-sibling offspring. These findings are of central importance for developing pedigree-based breeding programs for *H. illucens*. Although *H. illucens* females are known to lay unfertilized egg clutches (Dickerson *et al*., 2024; Harjoko *et al*., 2023; Tomberlin and Sheppard, 2002), it occurred at a low level in our experiment. This was the case even in the absence of males, which together with the high proportion of egg clutches that produced larvae shows that oviposition was a good but not perfect indicator of mating in our experiment.

Earlier studies have suggested that monogamy (Giunti *et al*., 2018) and polyandry (Hoffmann *et al*., 2021; Kotzé *et al*., 2019; Tomberlin and Sheppard, 2002) are the prevailing mating systems in *H. illucens*. Such mating systems would diminish the use of standard quantitative genetic designs, i.e. full-/half-sibling designs, because they require that males can successfully mate multiple females. We showed that polygyny occurs in laboratory reared *H. illucens*, supporting recent evidence of polygyny in mass breeding *H. illucens* populations (Chiabotto *et al*., 2024; Hoffmann *et al*., 2021; Jones and Tomberlin, 2021; Muraro *et al*., 2024). Our demonstration that polygyny can occur readily in a small container with only one single male facilitates the planning and development of full-/half-sibling designs in breeding programs for *H. illucens*. Full-/half-sibling breeding designs were recently used in house flies (Boatta *et al*., 2023; Hansen *et al*., 2024a), supporting our suggestion that advanced breeding programs are applicable in insects produced for food and feed. The findings of the current study pave the way for efficient breeding programs aiming at genetic improvement of traits of interest in *H. illucens*.

The shorter time until first mating when four females were present to a male instead of only one may have been caused by females disturbing each other, inducing movement and flight, and thereby increasing availability to the male. In theory, presenting multiple females to a single male also increases the chance that one of the females finds the male acceptable for mating due to complementary male genes or lower threshold for male quality (Hunt and Sakaluk, 2014; Moore and Moore, 1988). In *H. illucens*, male flight display and wing fanning are among traits assessed by females (Giunti *et al*., 2018; Julita *et al*., 2020). Mating is often highly dependent on female choice (Hunt and Sakaluk, 2014; Thornhill and Alcock, 1983), and female choice also plays a role in *H. illucens* when deciding whether to accept or reject a mating partner (Julita *et al*., 2020). Adding multiple females to a male will theoretically select for females that are more willing to mate under the given artificial conditions, which is a desired property in a production. This selection can also occur when only one female is present, as the male may not achieve mating the female. Our results showed no evidence that male fertility declines following one or multiple matings, and thus we saw no sign of sperm depletion. At the third and fourth consecutive mating, a presumed lower sperm cell concentration due to sperm depletion might have been counterbalanced by the generally longer time until next mating (Figure 3A) as well as the generally longer mating duration (Figure 3B), which might have aided to respectively produce and allocate more sperm (Manas *et al*., 2024). Very limited female remating was observed in the current study, evident through the high correlation between number of matings observed for a male and number of females laying egg clutches producing offspring sired by the same male. Although polyandry is demonstrated in *H. illucens* (Chiabotto *et al*., 2024; Muraro *et al*., 2024), female remating occurred to a very limited extent in our study as similarly reported in Giunti *et al*. (2018). In our setup, females did not have access to multiple males which could have stimulated the motivation for polyandry by broadening the sperm gene pool (Hunt and Sakaluk, 2014). It should be stated that we did not follow the offspring, and potential effects on offspring performance can therefore not be concluded upon.

In our study, LED lights were highly sufficient at inducing mating, supporting earlier studies (Heussler *et al*., 2018; Hoc *et al*., 2019; Julita *et al*., 2020; Liu *et al*., 2020; Macavei *et al*., 2020; Oonincx *et al*., 2016). Adaptation to artificial lighting instead of sunlight could potentially have occurred during rearing in captivity (Barrett *et al*., 2023). This is also the case for the lack of requirement for lekking, as defending lekking sites is typically not attainable at the high densities under production conditions as opposed to situations in nature (Tomberlin and Sheppard, 2001). Since our *H. illucens* culture had undergone several generations under commercial rearing, adaptation by natural selection to the production environment may have already occurred, and the flies from our investigated population might therefore be more willing to mate under artificial conditions than recently wild-caught flies. Since the populations kept by companies are largely highly commercialized and the population that we used originates from within the European commercialized clutch (Kaya *et al*., 2021), our findings will be relevant to most companies and institutions interested in breeding and genetically improving their *H. illucens* populations.

In conclusion, our demonstration that a selected male *H. illucens* can be mated individually with one or multiple virgin females and produce viable offspring with these shows that the reproductive biology of *H. illucens* allows for using quantitative genetic designs in breeding programs such as full-/half-sibling designs. The knowledge generated from such designs, i.e. heritabilities, genetic correlations and variance component estimates for traits of interest, are important for designing sustainable and efficient breeding programs. Using pedigree-based selection programs it is possible to control inbreeding rates, facilitate selection on multiple traits simultaneously, and sustain cumulative responses to selection across generations. However, relaxing sexual selection which is a consequence of the proposed mating scheme might have fitness consequences (Promislow *et al*., 1998), and potential risks related to this should be investigated in future studies.

## Acknowledgements

We thank H.J. MacLean, M.L. Schøn, I.E. Berggreen, and J.G. Sørensen for discussions and use of laboratory equipment. We also thank two anonymous reviewers whose critical assessments helped to improve our manuscript.

## References

Abdelmegeed, S.M., 2015. Effect of mating duration and the number of females/male moth of Bombyx mori L. on eggs fertility. Annals of Agricultural Sciences 60: 341–343. 10.1016/j.aoas.2015.10.008

Alcock, J., 1990. A large male competitive advantage in a lekking fly, Hermetia comstocki Williston (Diptera: Stratiomyidae). Psyche 97: 267–279. 10.1155/1990/72328

Athanassiou, C.G., Coudron, C.L., Deruytter, D., Rumbos, C.I., Gasco, L., Gai, F., Sandrock, C., De Smet, J., Tettamanti, G., Francis, A., Petrusan, J.-I. and Smetana, S., 2025. A decade of advances in black soldier fly research: from genetics to sustainability. Journal of Insects as Food and Feed 11: 219–246. 10.1163/23524588-00001122

Barrett, M., Chia, S.Y., Fischer, B. and Tomberlin, J.K., 2023. Welfare considerations for farming black soldier flies, Hermetia illucens (Diptera: Stratiomyidae): a model for the insects as food and feed industry. Journal of Insects as Food and Feed 9: 119–148. 10.3920/JIFF2022.0041

Boatta, F., Smit, J., Lautenschutz, M., Ellen, E. and Ellers, J., 2023. Heritability of fat accumulation in the house fly and its implication for the selection of nutritionally tailored phenotypes. Journal of Insects as Food and Feed 10: 825–834. 10.1163/23524588-20230149

Bunning, H., Rapkin, J., Belcher, L., Archer, C.R., Jensen, K. and Hunt, J., 2015. Protein and carbohydrate intake influence sperm number and fertility in male cockroaches, but not sperm viability. Proceedings of the Royal Society B 282: 20142144. 10.1098/rspb.2014.2144

Burdfield-Steel, E.R. and Harari, A.R., 2021. Special issue: ecology of sex and sexual communication in insects. Insects 12: 137. 10.3390/insects12020137

Cai, Z., Hansen, L.S., Laursen, S.F., Nielsen, H.M., Bahrndorff, S., Tomberlin, J.K., Kristensen, T.N., Sørensen, J.G. and Sahana, G., 2024. Whole-genome sequencing of two captive black soldier fly populations: implications for commercial production. Genomics 116: 110891. 10.1016/j.ygeno.2024.110891

Chiabotto, C., Grosso, F., Doretto, A. and Meneguz, M., 2024. Observation of mating behavior using marked flies of black soldier fly (Hermetia illucens) under sunlight condition. Journal of Insects as Food and Feed 10: 2017–2029. 10.1163/23524588-20230165

Clark, S.A., Hickey, J.M., Daetwyler, H.D. and van der Werf, J.H.J., 2012. The importance of information on relatives for the prediction of genomic breeding values and the implications for the makeup of reference data sets in livestock breeding schemes. Genetics Selection Evolution 44: 4. 10.1186/1297-9686-44-4

Cole, J.B., Dürr, J.W. and Nicolazzi, E.L., 2021. Invited review: the future of selection decisions and breeding programs: what are we breeding for, and who decides? Journal of Dairy Science 104: 5111–5124. 10.3168/jds.2020-19777

Dickerson, A.J., Lemke, N.B., Li, C. and Tomberlin, J.K., 2024. Impact of age on the reproductive output of Hermetia illucens (Diptera: Stratiomyidae). Journal of Economic Entomology 117: 1225–1234. 10.1093/jee/toae107

Dobermann, D., Swift, J. and Field, L., 2017. Opportunities and hurdles of edible insects for food and feed. Nutrition Bulletin 42: 293–308. 10.1111/nbu.12291

Emmerson, D.A., 1997. Commercial approaches to genetic selection for growth and feed conversion in domestic poultry. Poultry Science 76: 1121–1125. 10.1093/ps/76.8.1121

Eriksson, T. and Picard, C.J., 2021. Genetic and genomic selection in insects as food and feed. Journal of Insects as Food and Feed 7: 661–682. 10.3920/JIFF2020.0097

Facchini, E., Shrestha, K., van den Boer, E., Junes, P., Sader, G., Peeters, K., Schmitt, E., 2022. Long-term artificial selection for increased larval body weight of Hermetia illucens in industrial settings. Frontiers in Genetics 13: 1503. 10.3389/fgene.2022.865490

Falconer, D.S. and Mackay, T.F.C., 1996. Introduction to quantitative genetics. 4th edition. Addison Wesley Longman, Harlow, UK.

Fiske, P., Rintamäki, P.T. and Karvonen, E., 1998. Mating success in lekking males: a meta-analysis. Behavioral Ecology 9: 328–338. 10.1093/beheco/9.4.328

Giunti, G., Campolo, O., Laudani, F. and Palmeri, V., 2018. Male courtship behaviour and potential for female mate choice in the black soldier fly Hermetia illucens L. (Diptera: Stratiomyidae). Entomologia Generalis 38: 29–46. 10.1127/entomologia/2018/0657

Gligorescu, A., Chen, L., Jensen, K., Moghadam, N.N., Kristensen, T.N. and Sørensen, J.G., 2023. Rapid evolutionary adaptation to diet composition in the black soldier fly (Hermetia illucens). Insects 14: 821. 10.3390/insects14100821

Gowda, K.B., Jerry, D.R. and Zenger, K.R., 2025. Genetic improvement of farmed insect species: programmes, progress, and prospects. Journal of Insects as Food and Feed in press. 10.1163/23524588-00001387

Hansen, L.S., Laursen, S.F., Bahrndorff, S., Kargo, M., Sørensen, J.G., Sahana, G., Nielsen, H.M. and Kristensen, T.N., 2024a. Estimation of genetic parameters for the implementation of selective breeding in commercial insect production. Genetics Selection Evolution 56: 21. 10.1186/s12711-024-00894-7

Hansen, L.S., Laursen, S.F., Bahrndorff, S., Sørensen, J.G., Sahana, G., Kristensen, T.N. and Nielsen, H.M., 2024b. The unpaved road towards efficient selective breeding in insects for food and feed - A review. Entomologia Experimentalis et Applicata in press. 10.1111/eea.13526

Harjoko, D.N., Hua, Q.Q.H., Toh, E.M.C., Goh, C.Y.J. and Poniamoorthy, N., 2023. A window into fly sex: mating increases female but reduces male longevity in black soldier flies. Animal Behaviour 200: 25–36. 10.1016/j.anbehav.2023.03.007

Hawkes, M., Lane, S.M., Rapkin, J., Jensen, K., House, C.M., Sakaluk, S.K. and Hunt, J., 2022. Intralocus sexual conflict over optimal nutrient intake and the evolution of sex differences in life span and reproduction. Functional Ecology 36: 865–881. 10.1111/1365-2435.13995

Heussler, C.D., Walter, A., Oberkofler, H., Insam, H., Arthofer, W., Schlick-Steiner, B.C. and Steiner, F.M., 2018. Influence of three artificial light sources on oviposition and half-life of the black soldier fly, Hermetia illucens (Diptera: Stratiomyidae): improving small-scale indoor rearing. PLOS ONE 13: e0197896. 10.1371/journal.pone.0197896

Hill, W.G. and Kirkpatrick, M., 2010. What animal breeding has taught us about evolution. Annual Review of Ecology, Evolution, and Systematics 41: 1–19. 10.1146/annurev-ecolsys-102209-144728

Hoc, B., Noël, G., Carpentier, J., Francis, F. and Megido, R.C., 2019. Optimization of black soldier fly (Hermetia illucens) artificial reproduction. PLOS ONE 14: e0216160. 10.1371/journal.pone.0216160

Hoffmann, L., Hull, K.L., Bierman, A., Badenhorst, R., Bester-van der Merwe, A.E. and Rhode, C., 2021. Patterns of genetic diversity and mating systems in a mass-reared black soldier fly colony. Insects 12: 480. 10.3390/insects12060480

Hunt, J. and Sakaluk, S.K., 2014. Mate choice. In: Shuker, D.M. and Simmons, L. (eds.) The evolution of insect mating systems. Oxford University Press, Oxford, UK, pp. 129–158. 10.1093/acprof:oso/9780199678020.003.0008

Jensen, K., Kristensen, T.N., Heckmann, L.-H.L. and Sørensen, J.G., 2017. Breeding and maintaining high-quality insects. In: van Huis, A. and Tomberlin, J.K. (eds.) Insects as food and feed: from production to consumption. Wageningen Academic Publishers, Wageningen, the Netherlands, pp. 174–198. 10.3920/978-90-8686-849-0

Jensen, K., McClure, C., Priest, N.K. and Hunt, J., 2015. Sex-specific effects of protein and carbohydrate intake on reproduction but not lifespan in Drosophila melanogaster. Aging Cell 14: 605–615. 10.1111/acel.12333

Jensen, K. and Silverman, J., 2018. Frequently mated males have higher protein preference in German cockroaches. Behavioral Ecology 29: 1453–1461. 10.1093/beheco/ary104

Jones, B.M. and Tomberlin, J.K., 2021. Effects of adult body size on mating success of the black soldier fly, Hermetia illucens (L.) (Diptera: Stratiomyidae). Journal of Insects as Food and Feed 7: 5–20. 10.3920/JIFF2020.0001

Julita, U., Fitri, L.L., Putra, R.E. and Permana, A.D., 2020. Mating success and reproductive behavior of black soldier fly Hermetia illucens L. (Diptera, Stratiomyidae) in tropics. Journal of Entomology 17: 117–127. 10.3923/je.2020.117.127

Kaya, C., Generalovic, T.N., Ståhls, G., Hauser, M., Samayoa, A.C., Nunes-Silva, C.G., Roxburgh, H., Wohlfahrt, J., Ewusie, E.A., Kenis, M., Hanboonsong, Y., Orozco, J., Carrejo, N., Nakamura, S., Gasco, L., Rojo, S., Tanga, C.M., Meier, R., Rhode, C., Picard, C.J., Jiggins, C.D., Leiber, F., Tomberlin, J.K., Hasselmann, M., Blackenhorn, W.U., Kapun, M. and Sandrock, C., 2021. Global population genetic structure and demographic trajectories of the black soldier fly, Hermetia illucens. BMC Biology 19: 94. 10.1186/s12915-021-01029-w

Kortsmit, Y., van der Bruggen, M., Wertheim, B., Dicke, M., Beukeboom, L.W. and van Loon, J.A.A., 2023. Behaviour of two fly species reared for livestock feed: optimising production and insect welfare. Journal of Insects as Food and Feed 9: 149–169. 10.3920/JIFF2021.0214

Kotzé, R.C.M., Muller, N., du Plessis, L. and van der Horst, G., 2019. The importance of insect sperm: sperm ultrastructure of Hermetia illucens (black soldier fly). Tissue and Cell 59: 44–50. 10.1016/j.tice.2019.06.002

Lasley, J.F., 1978. Genetics of livestock improvement. 3rd edition. Prentice-Hall, Englewood, New Jersey, USA.

Laudani, F., Campolo, O., Latella, I., Modafferi, A., Palmeri, V. and Giunti, G., 2024. Does Hermetia illucens recognize sibling mates to avoid inbreeding depression? Entomologia Generalis 44: 1225–1232. 10.1127/entomologia/2024/2746

Laursen, S.F., Hansen, L.S., Bahrndorff, S., Nielsen, H.M., Sahana, G., Sørensen, J.G., Ørsted, M. and Kristensen, T.N., 2024. Genotype-by-environment interactions for mean performance and trait variation in house fly larvae reared on two diets. Entomologia Experimentalis et Applicata in press. 10.1111/eea.13530

Lemke, N.B., Dickerson, A.J. and Tomberlin, J.K., 2023. No neonates without adults: a review of adult black soldier fly biology, Hermetia illucens (Diptera: Stratiomyidae). BioEssays 45: 2200162. 10.1002/bies.202200162

Liceaga, A.M., 2021. Processing insects for use in the food and feed industry. Current Opinion in Insect Science 48: 32–36. 10.1016/j.cois.2021.08.002

Liu, Z., Najar-Rodriguez, A.J., Minor, M.A., Hedderley, D.I. and Morel, P.C.H., 2020. Mating success of the black soldier fly, Hermetia illucens (Diptera: Stratiomyidae), under four artificial light sources. Journal of Photochemistry and Photobiology B 205: 111815. 10.1016/j.jphotobiol.2020.111815

Lloyd, J.E., 1979. Mating behavior and natural selection. Florida Entomologist 62: 17–34. 10.2307/3494039

Lush, J.L., 1937. Animal breeding plans. Collegiate Press, Ames, Iowa, USA.

Macavei, L.I., Benassi, G., Stoian, V. and Maistrello, L., 2020. Optimization of Hermetia illucens (L.) egg laying under different nutrition and light conditions. PLOS ONE 15: e0232144. 10.1371/journal.pone.0232144

Manas, F., Piterois, H., Labrousse, C., Beaugeard, L., Uzbekov, R. and Bressac, C., 2024. Gone but not forgotten: dynamics of sperm storage and potential ejaculate digestion in the black soldier fly Hermetia illucens. Royal Society Open Science 11: 241205. 10.1098/rsos.241205

Meneguz, M., Miranda, C.D., Cammack, J.A. and Tomberlin, J.K., 2023. Adult behaviour as the next frontier for optimising industrial production of the black soldier fly Hermetia illucens (L.) (Diptera: Stratiomyidae). Journal of Insects as Food and Feed 9: 399–414. 10.3920/JIFF2022.0055

Meuwissen, T., 1997. Maximizing the response of selection with a predefined rate of inbreeding. Journal of Animal Science 75: 934–940. 10.2527/1997.754934x

Montanari, F., de Moura, A.P. and Cunha, L.M., 2021. Production and commercialization of insects as food and feed - Identification of the main constraints in the European Union. Springer, Cham, Switzerland. 10.1007/978-3-030-68406-8

Moore, A.J. and Moore, P.J., 1988. Female strategy during mate choice: threshold assessment. Evolution 42: 387–391. 10.2307/2409241

Morales-Ramos, J.A., Kelstrup, H.C., Rojas, M.G. and Emery, V., 2019. Body mass increase induced by eight years of artificial selection in the yellow mealworm (Coleoptera: Tenebrionidae) and life history trade-offs. Journal of Insect Science 19: 4. 10.1093/jisesa/iey110

Muraro, T., Lalanne, L., Pelozuelo, L. and Calas-List, D., 2024. Mating and oviposition of a breeding strain of black soldier fly Hermetia illucens (Diptera: Stratiomyidae): polygynandry and multiple egg-laying. Journal of Insects as Food and Feed 10: 1423–1435. 10.1163/23524588-20220175

Olivadese, M. and Dindo, M.L., 2023. Edible insects: a historical and cultural perspective on entomophagy with a focus on Western societies. Insects 14: 690. 10.3390/insects14080690

Oonincx, D.G.A.B., van Broekhoven, S., van Huis, A. and van Loon, J.J.A., 2015. Feed conversion, survival and development, and composition of four insect species on diets composed of food by-products. PLOS ONE 10: e0144601. 10.1371/journal.pone.0144601

Oonincx, D.G.A.B., Volk, N., Diehl, J.J.E., van Loon, J.J.A. and Belušič, G., 2016. Photoreceptor spectral sensitivity of the compound eyes of black soldier fly (Hermetia illucens) informing the design of LED-based illumination to enhance indoor reproduction. Journal of Insect Physiology 95: 133–139. 10.1016/j.jinsphys.2016.10.006

Perez-Staples, D., Prabhu, V. and Taylor, P.W., 2007. Post-teneral protein feeding enhances sexual performance of Queensland fruit flies. Physiological Entomology 32: 225–232. 10.1111/j.1365-3032.2007.00568.x

Permana, A.D., Fitri, L.L. and Julita, U., 2020. Influence of mates virginity on black soldier fly, Hermetia illucens L. mating performance, egg production and quality. Jurnal Biodjati 5: 174–181. 10.15575/biodjati.v5i2.9049

Promislow, D.E.L., Smith, E.A. and Pearse, L., 1998. Adult fitness consequences of sexual selection in Drosophila melanogaster. Proceedings of the National Academy of Sciences of the USA 95: 10687–10692. 10.1073/pnas.95.18.10687

Rapkin, J., Archer, C.R., Grant, C.E., Jensen, K., House, C.M., Wilson, A.J. and Hunt, J., 2017. Little evidence for intralocus sexual conflict over the optimal intake of nutrients for life span and reproduction in the black field cricket Teleogryllus commodus. Evolution 71: 2159–2177. 10.1111/evo.13299

Sarkar, K., Mandal, M. and Moorthy, S.M., 2009. Effect of mating duration and multiple use of male moth on reproductive performance of some cross breeds of silkworm, Bombyx mori L. International Journal of Industrial Entomology 19: 215–219.

Schneider, J.C., 2020. Effects of light intensity on mating of the black soldier fly (Hermetia illucens, Diptera: Stratiomyidae). Journal of Insects as Food and Feed 6: 111–119. 10.3920/JIFF2019.0003

Schultz, B., Serão, N. and Ross, J.W., 2020. Chapter 23 - Genetic improvement of livestock, from conventional breeding to biotechnological approaches. In: Bazer, F.W., Lamb, G.C. and Wu, G. (eds.) Animal Agriculture. Academic Press, London, UK, pp. 393–405. 10.1016/B978-0-12-817052-6.00023-9

Sellem, E., Paul, K., Donkpegan, A., Li, Q., Masseron, A., Chauveau, A., Gagnepain-Germain, F. and Lefebvre, T., 2024. Multitrait genetic parameter estimates in a Tenebrio molitor reference population: high potential for breeding gains. Animal 18: 101197. 10.1016/j.animal.2024.101197

Slagboom, M., Nielsen, H.M., Kargo, M., Henryon, M. and Hansen, L.S., 2024. The effect of phenotyping, adult selection, and mating strategies on genetic gain and rate of inbreeding in black soldier fly breeding programs. Genetics Selection Evolution 56: 71. 10.1186/s12711-024-00938-y

Thornhill, R. and Alcock, J., 1983. The evolution of insect mating systems. Harvard University Press, Cambridge, Massachusetts, USA. 10.4159/harvard.9780674433960

Tomberlin, J.K. and Sheppard, D.C., 2001. Lekking behavior of the black soldier fly (Diptera: Stratiomyidae). Florida Entomologist 84: 729–729. 10.2307/3496413

Tomberlin, J.K. and Sheppard, D.C., 2002. Factors influencing mating and oviposition of black soldier flies (Diptera: Stratiomyidae) in a colony. Journal of Entomological Science 37: 345–352. 10.18474/0749-8004-37.4.345

Tomberlin, J.K. and van Huis, A., 2020. Black soldier fly from pest to ‘crown jewel’ of the insects as feed industry: an historical perspective. Journal of Insects as Food and Feed 6: 1–4. 10.3920/JIFF2020.0003

van Huis, A., 2020. Insects as food and feed, a new emerging agricultural sector: a review. Journal of Insects as Food and Feed 6: 27–44. 10.3920/JIFF2019.0017

van Huis, A., 2021. Prospects of insects as food and feed. Organic Agriculture 11: 301–308. 10.1007/s13165-020-00290-7

van Huis, A., Oonincx, D.G.A.B., Rojo, S. and Tomberlin, J.K., 2020. Insects as feed: house fly or black soldier fly? Journal of Insects as Food and Feed 6: 221–229. 10.3920/JIFF2020.x003

Worden, B.D. and Parker, P.G., 2001. Polyandry in grain beetles, Tenebrio molitor, leads to greater reproductive success: material or genetic benefits? Behavioral Ecology 12: 761–767. 10.1093/beheco/12.6.761

Zaalberg, R.M., Nielsen, H.M., Noer, N.K., Schou, T.M., Jensen, K., Thormose, S., Kargo, M. and Slagboom, M., 2024. A bio-economic model for estimating economic values of important production traits in the black soldier fly (Hermetia illucens). Journal of Insects as Food and Feed 10: 1411–1421. 10.1163/23524588-00001126

